# JAFFAL: Detecting fusion genes with long read transcriptome sequencing

**DOI:** 10.1101/2021.04.26.441398

**Authors:** Nadia M. Davidson, Ying Chen, Teresa Sadras, Georgina L. Ryland, Piers Blombery, Paul G. Ekert, Jonathan Göke, Alicia Oshlack

## Abstract

Massively parallel short read transcriptome sequencing has greatly expanded our knowledge of fusion genes which are drivers of tumor initiation and progression. In cancer, many fusions are also important diagnostic markers and targets for therapy. Long read transcriptome sequencing allows the full length of fusion transcripts to be discovered, however, this data has a high rate of errors and fusion finding algorithms designed for short reads do not work. While numerous fusion finding algorithms now exist for short read RNA sequencing data, there are few methods to detect fusions using third generation or long read sequencing data. Fusion finding in long read sequencing will allow the discovery of the full isoform structure of fusion genes.

Here we present JAFFAL, a method to identify fusions from long-read transcriptome sequencing. We validated JAFFAL using simulation, cell line and patient data from Nanopore and PacBio. We show that fusions can be accurately detected in long read data with JAFFAL, providing better accuracy than other long read fusion finders and with similar performance as state-of-the-art methods applied to short read data. By comparing Nanopore transcriptome sequencing protocols we find that numerous chimeric molecules are generated during cDNA library preparation that are absent when RNA is sequenced directly. We demonstrate that JAFFAL enables fusions to be detected at the level of individual cells, when applied to long read single cell sequencing. Moreover, we demonstrate JAFFAL can identify fusions spanning three genes, highlighting the utility of long reads to characterise the transcriptional products of complex structural rearrangements with unprecedented resolution. JAFFAL is open source and available as part of the JAFFA package at https://github.com/Oshlack/JAFFA/wiki.

## Background

Genomic rearrangements are common in the landscape of cancer and when breakpoints occur within different genes these can be transcribed into a new hybrid transcript, producing a so-called fusion gene. Fusions may drive cancer through activation of onocogenes [1] or inactivation of tumour suppressors. Often such fusions are recurrent across patient cohorts and novel drugs have been developed to specifically target a number of them [2]. Fusion detection can therefore inform cancer care, and eliciting their function in cancer initiation and progression is an ongoing area of research.

Over the last decade, massively parallel short read transcriptome sequencing has greatly expanded our knowledge of fusion genes across cancers and is increasingly being used for clinical diagnostics [3–5]. For example, The Cancer Genome Atlas (TCGA) utilised short read transcriptome sequencing across a range of tumour types to estimate that approximately 16% of cancers have a fusion event which drives the disease [6]. Fusion discovery through sequencing has necessitated the development of dedicated bioinformatics methods. Since the advent of the first approaches [7,8], fusion finding has improved in both accuracy and speed, and there are now numerous tools available [9–12].

Third generation, or long read sequencing technologies, as offered by Oxford Nanopore Technologies (ONT) [13] and Pacific Bioscience (PacBio) [14], can provide novel insight into fusions and their role in cancer. Unlike short read sequencing, long read sequencing does not require fragmentation, hence the full length of individual mRNA molecules can be sequenced. Long range information about the structure and sequence of fusion transcripts, including splicing, SNPs or additional structural variants, not immediately adjacent to the breakpoint can be obtained. This offers to improve predictions of open reading frames, protein sequence and therefore biological relevance. Around 12% of fusions analysed by the Pan-Cancer Analysis of Whole Genomes (PCAWG) Consortium were supported by multiple genomic rearrangements [15]. Long read sequencing will allow us to understand how these complex structural changes are transcribed into RNA. Long read sequencing has several other advantages, for example ONT allows RNA to be sequenced directly, without reverse transcription and therefore RNA modifications can be measured [16]. In addition, rapid and remote diagnostics may be possible with ultra portable sequencing machines and rapid workflows [17,18]. Finally, new protocols allow full length sequencing of genes at the level of single cells [19–21].

Most fusion finders rely on short read alignment algorithms, which are incapable of accurately and efficiently mapping long reads [22]. An additional challenge is that the raw data generated by third generation technologies have a high rate of errors [23], in particular insertion and deletions, that short read algorithms were not designed to account for. As a result, to the best of our knowledge, only three fusion finding methods are available for long read transcriptome data: JAFFA [24] is a pipeline we previously developed and although it can process transcriptome sequencing data of any length, it has low sensitivity when error rates are high; Aeron [25] detects fusions by aligning long reads to a graph based representation of the reference transcriptome; and LongGF [26] analyses genome mapped long read data and detects fusions by identifying reads aligning to multiple genes. An additional program, NanoGF [26] can detect fusions in long read genome sequencing data, but is not designed for transcriptome sequencing.

To take advantage of new long read sequencing technologies for fusion finding and characterisation, we have developed JAFFAL, a new method which is built on the concepts developed in JAFFA and overcomes the high error rate in long read transcriptome data by using alignment methods and filtering heuristics which are designed to handle noisy long reads. We validated JAFFAL using simulated data as well as cancer and healthy cell line data for ONT and PacBio. By comparing ONT transcriptome sequencing protocols we show that numerous chimeric molecules are generated during cDNA library preparation that are absent when RNA is sequenced directly. JAFFAL effectively filtered these events by accurately determining break-point positions relative to exon boundaries. We show JAFFAL is the most accurate fusion finder available for noisy long read data, allowing fusions to be detected in long read data with similar accuracy as short reads. On two patient ONT sequencing samples, JAFFAL was able to detect clinically relevant fusions. Finally, as a proof-of-feasibility, we apply JAFFAL to long read single cell sequencing of five cancer cell lines and demonstrate its ability to recover known fusions at the level of individual cells. Furthermore, by utilising full length transcript information in the long reads we identified BMPR2-TYW5-ALS2CR11, a fusion composed of three genes, in individual cells of the H838 non-small-cell lung cancer cell line. JAFFAL is open source and available as part of the fusion finding package JAFFA, versions 2.0 and higher (https://github.com/Oshlack/JAFFA/wiki).

## Results and Discussion

### JAFFAL pipeline

JAFFAL is a new multistage pipeline (Figure 1) written in bpipe [27] and was motivated by our approach from the *Direct* mode of JAFFA [24]. The pipeline consists of the following steps: (1) Fusions are detected by first aligning long reads to a reference transcriptome (hg38 gencode version 22) [28] using the noise tolerant long read aligner minimap2 [29]. (2) Reads consistent with a fusion gene, ie. those with sections aligning to different genes, are selected for further analysis and (3) subsequently aligned to the reference genome hg38, also using minimap2. Reads which do not span multiple genes after reference genome alignment are removed. This double alignment, to a reference transcriptome and genome, ensures that false positives are minimised, and reduces computational time, as only a small subset of reads need to be aligned to the full reference genome.

**Figure 1.**
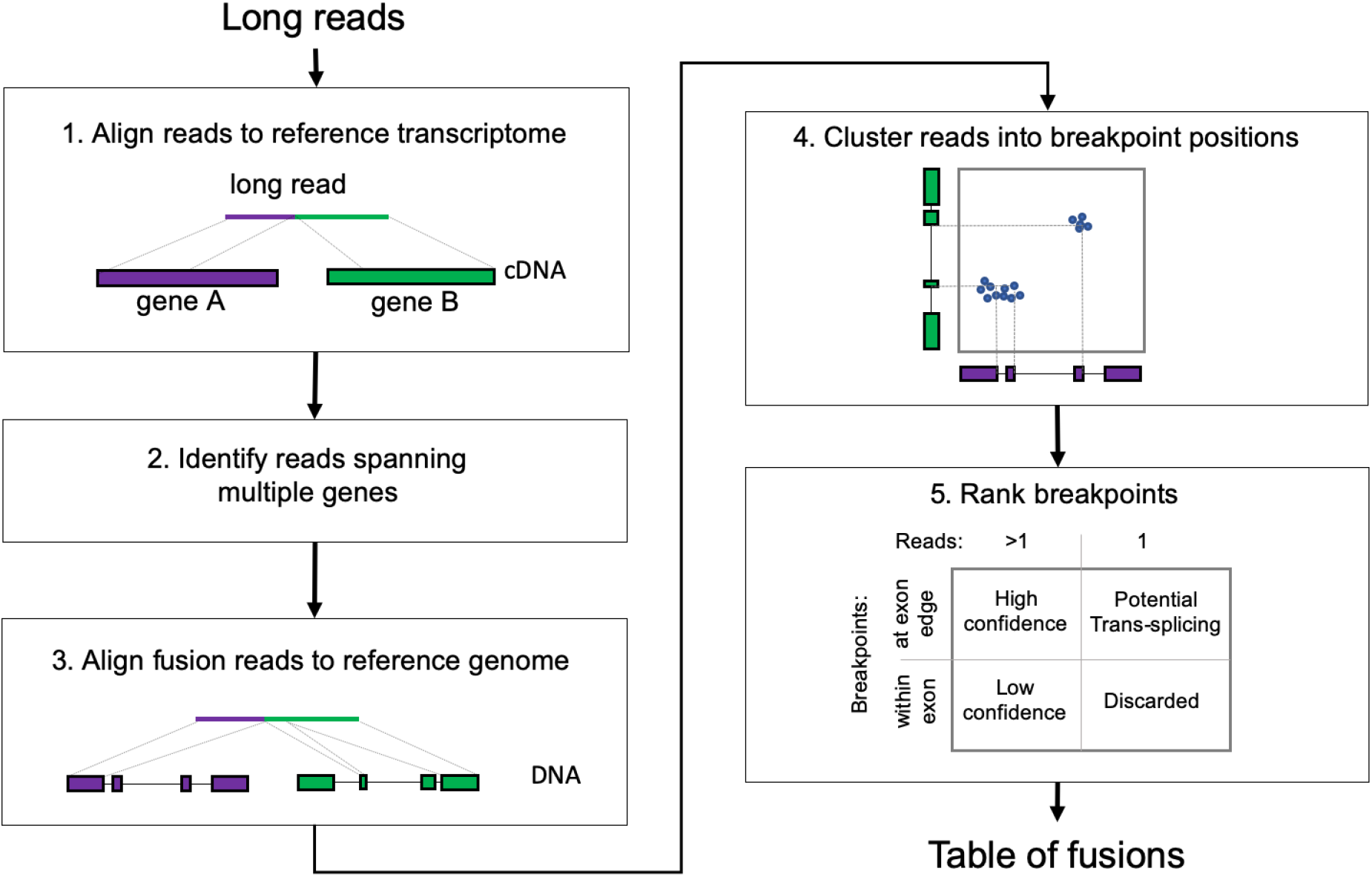
JAFFAL pipeline steps for fusion detection. Reads are aligned to the reference transcriptome, reads split across different genes are identified as candidate fusion reads and subsequently aligned to the reference genome for confirmation. Reads are clustered into breakpoint positions which are then ranked and reported (see text for details).

Next, (4) JAFFAL uses the end position of reference genome alignments to determine fusion breakpoints. Due to the high error rate in long read sequencing, alignment end positions may be inaccurate. To account for this, JAFFAL employs a strategy which anchors transcript breakpoints to exon boundaries. While structural rearrangements commonly occur within introns, splice sites are usually preserved, creating fusion transcripts where the breakpoint in the RNA is at the end or start of an exon. JAFFAL will realign breakpoints to the exon boundaries if exon boundaries are identified within 20bp of the original alignment breakpoints. This is only done if the adjustments on the 5’ and 3’ sides of the break are consistent with one another, and result in a new breakpoint at exon boundaries for both the 5’ and 3’ gene. All such exon boundary breakpoints will be reported by JAFFAL.

Due to insertion and deletion errors, or genuine breakpoints within an exon body, many reads will not satisfy the requirements for breakpoint adjustment. These reads are clustered by genomic position. One breakpoint is reported for each cluster, which will be either the one preserving exon boundaries, or the one with the highest read support. Clustering is achieved by iterating through all non-exon boundary breakpoints, starting with the one with the least read support. The breakpoint’s reads will be reassigned to the closest breakpoint from other reads within 50bp (euclidean genomic distance). If no other breakpoint is found within 50bp the breakpoint is reported.

Finally, (5) breakpoints are ranked into “High Confidence”, “Low Confidence” and “Potential Trans-Splicing” classes (Figure 1), similar in concept to the ranking in JAFFA for short reads [24]. “High Confidence” fusions are supported by two or more reads with breakpoints aligning to exon boundaries. “Low Confidence” fusions are also supported by two or more reads, but breakpoints do not align to exon boundaries. “Potential Trans-Splicing” events are supported by a single read, with breakpoints aligning to exon boundaries (Figure 1). Numerous “Potential Trans-Splicing” events are seen in healthy RNA-Seq samples [24,30], and should generally be filtered out. However some true fusions may be reported as “Potential Tran-Splicing”, for example those with low expression levels or in samples with low tumour purity. All other events are removed. Run-through transcription, identified by breakpoints within 200kbp of each other and where the genes are transcribed in the same order as the reference genome, are also filtered out by default, as are fusions which involve the mitochondrial chromosome. However, these events may be recovered by the user if needed.

For each breakpoint which passes filtering, JAFFAL reports the genes involved, genomic coordinates, number of reads supporting the event, ranking class, whether it is inframe and whether it has been seen before in the Mitelman database of genomic rearrangements [31]. Within each class, breakpoints are ranked by the number of supporting reads. Finally, rare multi-fusion events, which incorporate sequences from three or more genes, are identified by searching for reads with two or more breakpoints in the final list. These are reported in a separate table.

### Simulated fusions are accurately detected in noisy long read data with JAFFAL

JAFFAL’s ability to detect fusions was tested on simulated data for the same 2500 fusion events simulated by Haas et al. [10]. For each fusion, Hass et al. selected two protein-coding genes at random. The breakpoint within each fusion was decided by joining a randomly selected exon from each gene, requiring a minimum 100bp of sequence from each. We simulated long reads from the resulting fusion gene sequences using Badread version 0.1.5 [32], which uses a noise model based on real data. The 2500 fusions were divided into 25 groups with varying coverage and read identity levels. Specifically, 500 fusion events were simulated across 5 coverage levels: x1, x2, x10, x50 and x100 reads. For each coverage we simulated 100 fusions each with a mean read identity of 75%, 80%, 85%, 90% and 95% (standard deviation 5%). These read identities were designed to cover the range expected in real data. For example, the cell line data used to validate JAFFAL was estimated to have read identities in the range 80 to 85% for ONT and 85% to over 95% for PacBio (Supplementary Figure 1). Fusions were considered detected if a breakpoint was reported within 1kbp euclidean distance of the simulated breakpoint. Fusions were simulated with both ONT and PacBio noise models. To emulate a realistic background, we combined the simulated ONT reads with 25 million cDNA reads from the non-tumour reference cell line NA12878 generated by the Nanopore WGS consortium [33] where few fusions should be present. JAFFAL was found to have similar fusion finding sensitivity across the three datasets: ONT simulation without background, PacBio simulation without background and ONT simulation with background (Figure 2, Supplementary Figure 2).

**Figure 2:**
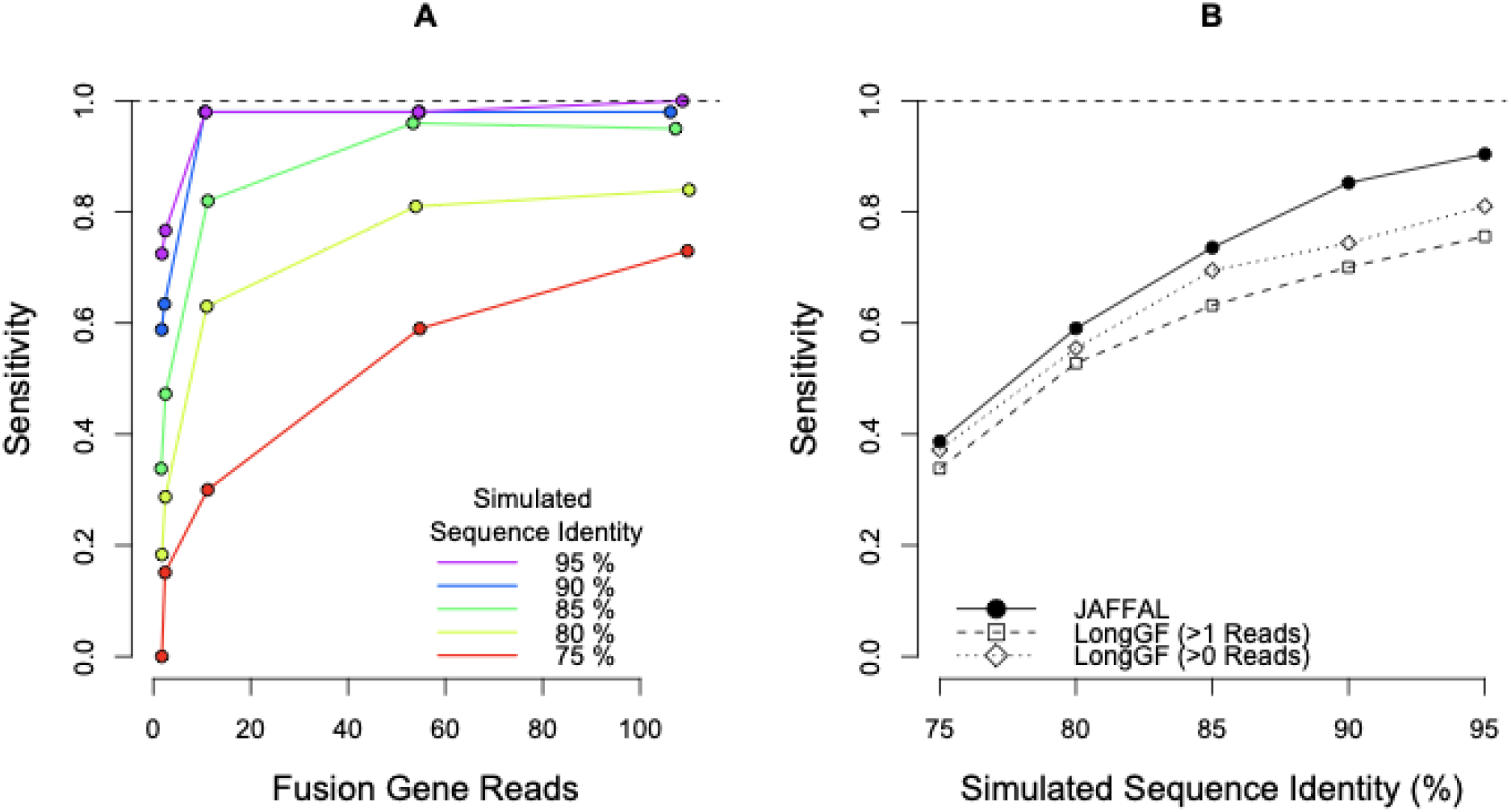
Fusion finding sensitivity on simulated ONT data with background. A) The fraction of simulated fusions detected (y-axis) by JAFFAL across a range of fusion coverage levels (x-axis). Read identity levels are shown in different colours (red-purple). B) The fraction of simulated fusions detected (y-axis) by JAFFAL and LongGF for sequence identity levels of 75-95%.

JAFFAL detected 98% of simulated fusions when the read identity was 90% or above and the coverage was 10 or greater (Figure 2A, Supplementary Figures 2A and 2C). Across all simulated fusions that were detected, approximately 84% were classed as high confidence and 99% with a single breakpoint. As expected, with low coverage and read identity, fewer fusions were detected. High error rates also impacted the fraction of supporting reads identified by JAFFAL. Amongst the fusions detected, the reported supporting reads were only 14% of the simulated coverage when the identity was 75%, compared to 79% of coverage when the identity was 95%. Most reads which failed to be reported did not align to two genes in the initial reference transcriptome mapping. This impacted up to 84% of simulated reads when the read identity was 75%, with 40%failing to align to even one gene. The number of reads lost in other stages of the JAFFAL pipeline remained low, approximately 10%, across all scenarios (Supplementary Figure 3).

JAFFAL’s sensitivity on the simulated data was comparable to the alternative long read fusion finder, LongGF’s when the data contained only fusion reads (Supplementary Figures 2B and 2D). However, in the presence of background reads from NA12878, JAFFAL had higher sensitivity than LongGF (Figure 2B), even after reducing LongGF’s default parameter of >1 read support to >0 read support. As we were unsuccessful in running Aeron, results for that program are not shown. We did not compare to NanoGF as it is designed for data from genome sequencing rather than whole transcriptome sequencing. JAFFAL was also found to have superior breakpoint resolution to LongGF; for 96% of fusions detected by JAFFAL, the exact breakpoint was reported, compared to just 2% from LongGF. However, almost all breakpoints were within 20bp of the simulated position for both tools.

### JAFFAL’s fusion ranking is effective at separating false positives in non-tumor cell line data

To assess the false positive rate of JAFFAL across different classification levels and sequencing protocols, we applied it to ONT direct RNA and amplified cDNA sequencing of NA12878 without simulated fusion events. Hence, almost all fusions reported should be false positives. For both protocols JAFFAL reported few fusions with a ranking of high confidence as expected (Table 1, Supplementary Table 1). Amongst the high confidence calls, three were common to both the direct RNA and cDNA datasets. One of these, KANSL1-ARL17A, is a germline fusion known to be present in a subset of the healthy population [34]. The two other fusions were consistent with run through transcription, where the distance between breakpoints just exceeded the 200kbp threshold for filtering. A further two fusions called in the cDNA sample could be explained as run through transcription for the same reason. JAFFAL reported several hundred “Potential Trans-splicing” events, which was consistent with levels seen previously from short read sequencing [24]. LongGF detected just five fusions with multi-read support for the direct RNA protocol, all of which were also reported by JAFFAL (two as high and three as low confidence).

**Table 1:** The number of fusion genes and breakpoints called in the non-cancer cell line NA12878 from ONT direct RNA and amplified cDNA. Most calls are presumed to be false positives. The number of fusions in the highest rank category for each tool is shown in bold. We hypothesize that most of the multi-read fusions reported by LongGF applied to the cDNA dataset (173) are chimeras introduced during library preparation. JAFFAL ranks these events as Low Confidence. The number of breakpoints for LongGF are not shown as it only reports one breakpoint per fusion gene by default.

|  |  | Direct RNA |  |  | cDNA |  |  |
| --- | --- | --- | --- | --- | --- | --- | --- |
| Total Reads Processed |  | 14,971,421 |  |  | 25,418,307 |  |  |
|  |  | Fusion Genes | Break Points | Reads Support: Median (Range) | Fusion Genes | BreaP Points | Reads Support: Median (Range) |
| Fusion genes called by JAFFAL | High Confidence | **4** | 4 | 4.5 (2-14) | **8** | 8 | 6 (2-24) |
|  | Low Confidence | 5 | 7 | 2 (2-11) | 94 | 121 | 2 (2-49) |
|  | Potential Trans-splicing | 344 | 344 | 1 (1-1) | 412 | 412 | 1 (1-1) |
| Fusion genes called by Long GF | > 1 Read support | **5** |  | 2 (2-14) | **173** |  | 2 (2-522) |
|  | = 1 Read support | 713 |  | 1 (1-1) | 386 |  | 1 (1-1) |

On the cDNA data, LongGF reported 173 fusions with multi-read support whereas JAFFAL only called 8 fusions as high confidence. The number of low confidence calls from JAFFAL was similar to LongGF however with direct RNA (5 fusions reported) compared to the cDNA protocol (94 fusions reported) (Table 1). We hypothesise this is due to chimeric molecule creation during cDNA library preparation [35,36]. These chimeras are distinct from the chimeras commonly seen in nanopore data due to ligation, where two full length transcripts are joined. The chimeras detected by JAFFAL do not contain an internal adapter sequence and only part of each gene is seen in the sequence (Supplementary Table 2). To ensure the excess in low confidence fusions in the cDNA sample was not a result of larger library size, we downsampled the reads to the direct RNA library size (Supplementary Table 3) and 43 low confidence fusions were still observed.

The ranking of these fusions as low confidence is consistent with the hypothesis that they are created during library preparation. A hallmark of these events are breakpoints occurring within exons, rather than at exon boundaries and allows them to be separated from true fusions by JAFFAL’s ranking. LongGF does not appear to separate this class of artifact and reported a large number of false positives in the cDNA dataset (Table 1).

To examine chimeras further, we switched off JAFFAL’s default filtering of mitochondrial genes, and looked at the prevalence of fusions reported between a gene on the mitochondrial chromosome and a gene on another chromosome. These are likely to be chimeras which are not native to cells. 116 such mitocondrial chimeras were reported by JAFFAL in the cDNA library at low confidence. None were reported at other confidence levels or in the direct RNA library. To confirm this result in an independent dataset, we examined chimeras in data from five cell lines from the Singapore Nanopore-Expression Project, SGNex [37], where several replicates of direct RNA, direct cDNA and amplified cDNA ONT sequencing are available. A much lower rate of mitochondrial chimeras was seen in the direct RNA sequencing, but no significant difference was observed between direct and amplified cDNA (Supplementary Figure 4).

These results demonstrate that chimeras created during library preparation can be effectively separated from true fusions if fusion breakpoints are accurately determined and their position relative to exon boundaries used. The absence of chimeras in direct RNA sequencing is striking and gives confidence in the fusions called from this protocol. In particular, the filtering based on breaks occurring at the exon boundary can be removed allowing confident detection of the rare instances where a breakpoint occurs within an exon.

### JAFFAL detects known fusions in cancer cell lines

To further confirm JAFFAL’s accuracy, it was applied to public long read transcriptome sequencing of six cancer cell line, where fusions had been previously validated using RT-PCR and Sanger sequencing, or there was orthogonal evidence of a translocation from whole genome sequencing [4,38–47] (Table 2, Supplementary Table 4). The four cell lines MCF-7, HCT-116, A549 and K562 were sequenced with ONT and are available as part of SGNex [37]. The direct RNA, direct cDNA and amplified cDNA replicates were combined into a single fastq file for fusion calling on each cell line. These samples had estimated read identities of 80-85% (Supplementary Figure 1). The three cell lines MCF-7, HCT-116 and SK-BR-3 [43] which had PacBio SMRT sequencing were downloaded from the Sequence Read Archive (SRA) and had estimated read identities over 95% (MCF-7 and HCT-116) and ~86% (SK-BR-3). Fusion genes reported by JAFFAL and LongGF were compared to those previously validated using gene identifiers. When a fusion had multiple breakpoints, we assigned the fusion gene the classification of its highest rank breakpoint.

**Table 2:**
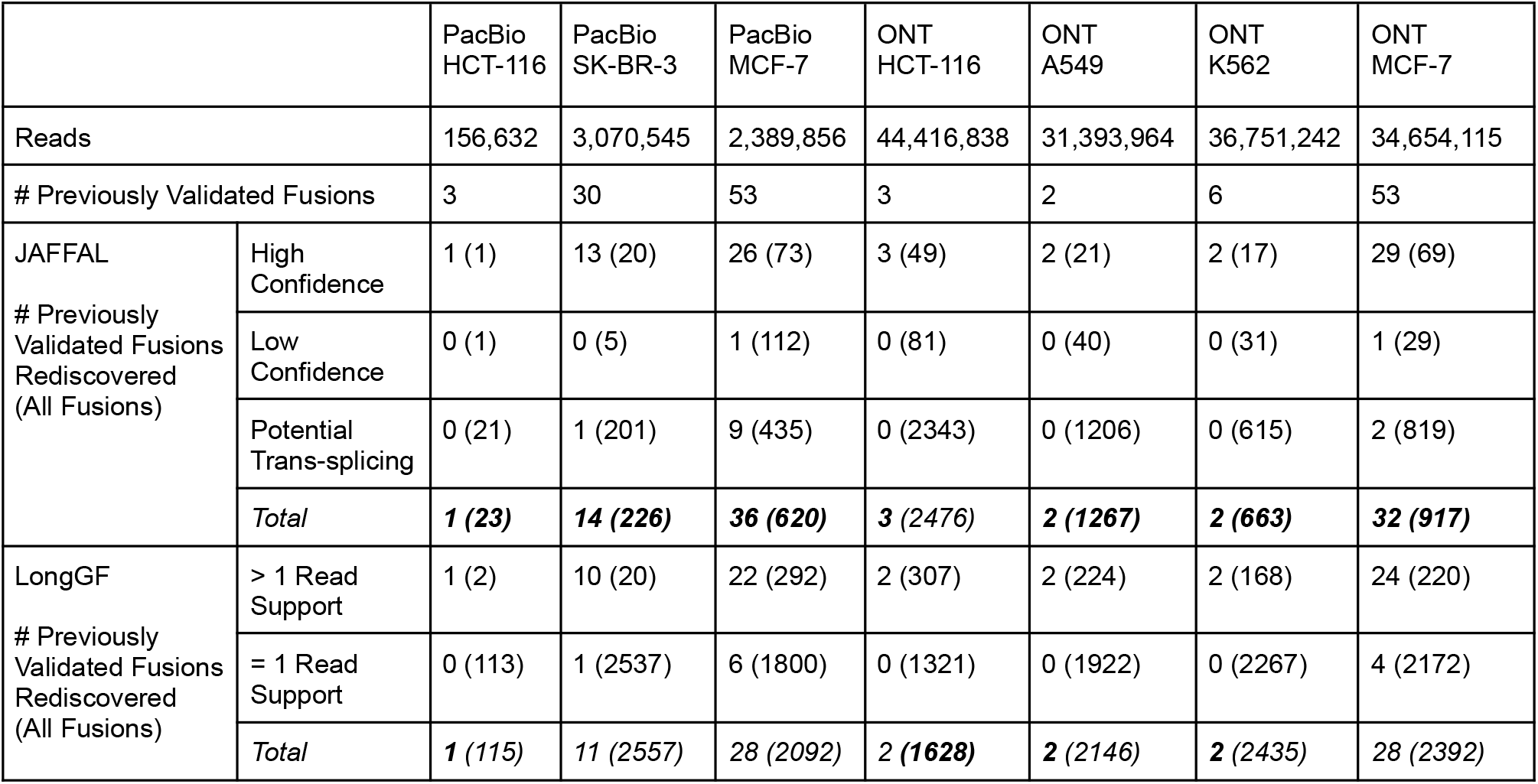
The number of previously validated fusions rediscovered across seven long read sequencing datasets by JAFFAL and LongGF. The total number of fusion genes reported by each tool, including those not previously validated, are indicated in parentheses.

JAFFAL rediscovered approximately half the previously validated fusion genes (Table 2) and 84% of these were ranked as high confidence. Previously validated fusions were reported with a range of supporting reads: 1-2,929 (median=15) and breakpoints 1-13 (median=1) (Supplementary Tables 5 and 6). Compared to LongGF, JAFFAL reported equal or more previously validated fusions for all datasets and ranked them higher (Figure 3A and B, Table 2). All fusions were reported with genes in the correct 5’ and 3’ order for JAFFAL compared to 68% for LongGF. JAFFAL also reported fewer total fusions in six of seven datasets, with unvalidated detections likely to be predominately false positives. These were reported by JAFFAL to be mainly in the potential trans-splicing category similar to those seen in the reference cell line NA12878.

**Figure 3:**
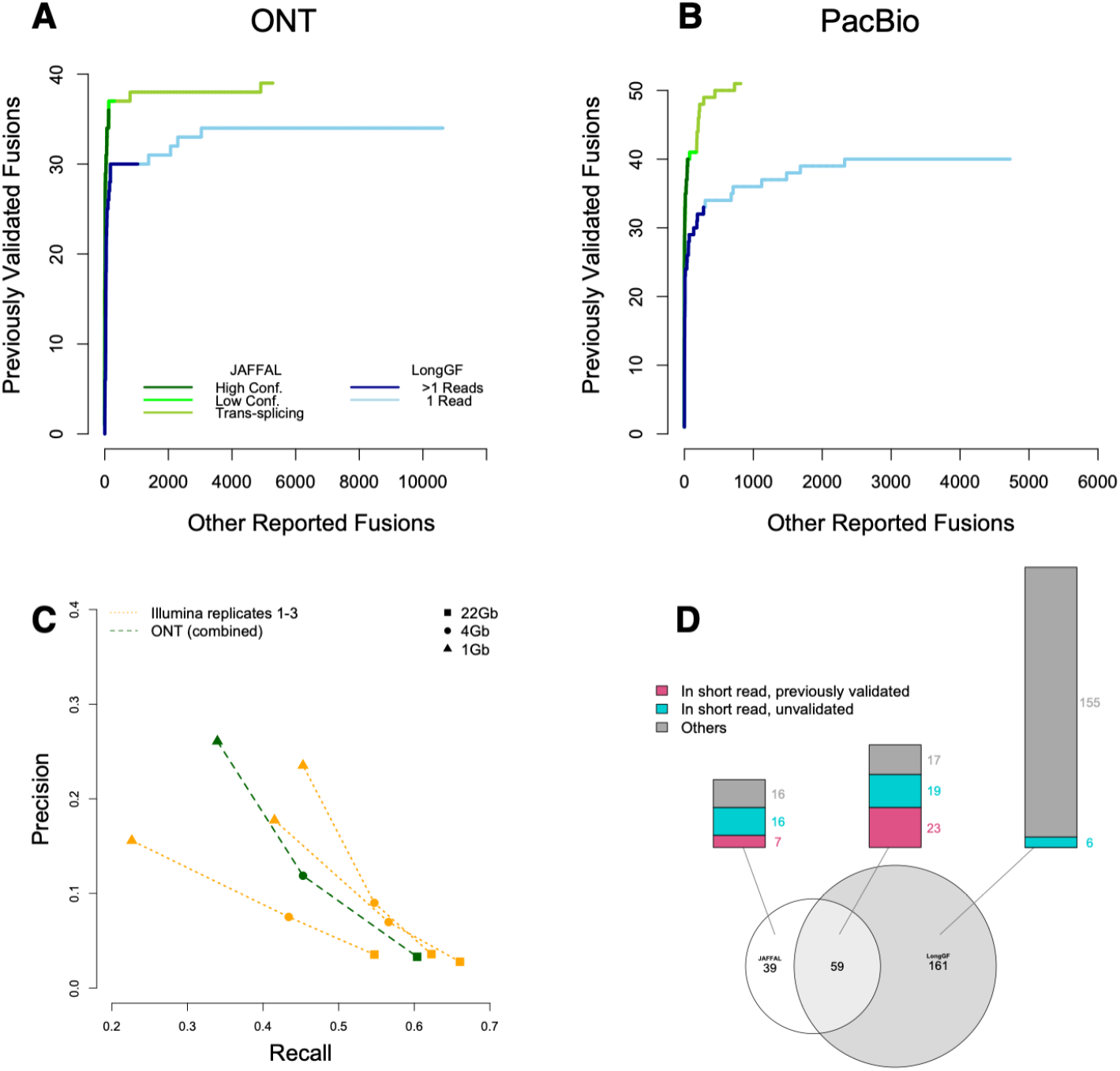
Comparison of JAFFAL and LongGF on cancer cell line sequencing. Shown are ROC style curve with the ranking of previously validated fusions against other reported fusions for A) MCF-7, HCT-116, A549 and K562 cell lines sequenced with ONT and B) MCF-7, HCT-116 and SK-BR-3 cell lines sequenced with PacBio. C) For MCF-7 only, fusions from JAFFAL are compared against three short read Illumina replicates across three sequencing depths. Precision and recall are calculated using previously validated fusions in MCF-7. D) The overlap between fusions called by JAFFAL (high and low confidence) and LongGF (> 1 read support) on MCF-7.

Although JAFFAL failed to report a number of the previously validated fusions, this could be caused by differences in sequencing depth and cell line batch effects, and is something that has been observed previously in short read data [24]. Hence, we also benchmarked JAFFAL’s sensitivity against fusions called on matched short read data from the same samples. We used MCF-7 from SGNex, as this cell line had the greatest number of validated fusions. Fusions were called in the short read data with JAFFA, which has been independently benchmarked in several studies [10,48,49]. Because the long and short read sequencing were from the same replicates, we would expect a similar set of fusions to be expressed. The short read data was significantly more deeply sequenced (137 million 150bp paired-end Illumina reads), hence we subsampled the datasets to approximately 22Gbp, 4Gbp, and 1Gbp to compare performance over a range of depths. The precision and recall of JAFFAL on long reads was found to be within the range of short read replicates, demonstrating both the accuracy of JAFFAL and the utility of noisy long read data for fusion detection more generally (Figure 3C).

The short read data of MCF-7 was also used to assess the likelihood of JAFFAL’s unvalidated calls being genuine fusions. Genuine fusions should be found in both the ONT and full depth Illumina data. Of the 69 high confidence fusion genes called by JAFFAL, 60 were also detected in the short read data. 5 of 29 low confidence and 19 of 819 potential trans-splicing events were common, indicating that events in these categories are more likely to be artifacts, such as chimeras generated during library preparation, which is consistent with results from NA12878.

Breakpoint positions from JAFFAL were also consistent with short read data. For the 84 fusion genes common to the short and long read data across all confidence levels, 140 different breakpoints were reported by JAFFAL (range: 1-13 per fusion pair) and 181 by JAFFA (range: 1-15 per fusion pair) on the short read data. 117 of these were common between the short and long read datasets (within 20bp), with the majority, 104, an exact match. Note that the number of breakpoints is greater than the number of fusion genes likely due to alternative splicing. Correctly identifying all breakpoints is important for determining whether any fusion transcript is in-frame.

Overall on the MCF-7 ONT cell line data, JAFFAL’s high and low confidence calls showed consistency with previously validated fusions, fusions in matched short read data and fusions called by LongGF (Figure 3D). Only 16% of fusion genes reported by JAFFAL as high or low confidence were not seen by other approaches, compared to 70% of LongGF’s calls (>1 read support). Taken together, these results suggest JAFFAL is highly accurate, in particular in the high confidence class.

### Detection of clinically relevant fusions with long read sequencing in leukemia

JAFFAL was next applied to two samples from patients with leukemia to assess its ability to detect fusions in a real-word context. One patient had acute myeloid leukemia (AML) with a RUNX1-RUNX1T1 fusion and cDNA sequencing was performed by Lui et al. [26] on ONT GridION, resulting in 8 million reads. The other patient had B-cell acute lymphoblastic leukemia (B-ALL) with the rare phenomenon of both BCR-ABL1 and IGH-CRLF2 fusions detected by cytogenetics and short read RNA sequencing. ONT sequencing was performed on amplified cDNA with a MinION, resulting in 13 million reads.

JAFFAL detected the RUNX1-RUNX1T1 and BCR-ABL1 fusions ranked as first of 17 and fifth of 51 high confidence calls in their respective samples. Consistent with results from simulation and cell line data, JAFFAL found the exact breakpoints. However it failed to detect the IGH-CRLF2 fusion, despite the fusion transcript being evident through manual inspection in the sequencing data. IGH-CRLF2 was missed because the breakpoint occurred approximately 2kbp upstream of CRLF2 and is an example of enhancer hijacking. Inability to detect fusions involving intergenic regions is an important limitation of JAFFAL, but is one shared by most fusion finders, with a few exceptions [9,50]. LongGF also failed to detect the IGH-CRLF2 fusion (Supplementary Table 7).

### Fusion detection at the single cell level

Single cell transcriptomics using long read sequencing is emerging as a powerful system to investigate transcript diversity across cell types [19–21]. As tumour samples nearly always contain multiple cell types, including infiltration of immune cells [51], or multiple clones [52], it is of broad interest to track the presence of fusion genes within single cells. As a proof of the feasibility for calling fusions at the single cell level, we applied JAFFAL to public data from a mixed sample of five cancer cell lines that was sequenced with ONT in combination with 10x Genomics and Illumina sequencing (Supplementary Table 8) [20]. A total of 18 million ONT reads could be assigned cellular barcodes across 557 cells. As expected, cells clustered into five distinct groups based on gene expression from short read data (Figure 4A). High confidence fusions called by JAFFAL and found in four or more cells were investigated further. JAFFAL identified 15 fusions, with a range of read support of 1-14 (median=1) per cell. Cells where fusions were identified had a range of 854-147,531 reads in library size (median=43,660) (Supplementary Figure 5). Of the fusions, 13 were also found in short read RNA-seq of the same cell lines as part of the Cancer Cell Line Encyclopedia, CCLE [53] (Figure 4B). Distinct sets of fusions were associated with each cluster, enabling the annotation of the cluster to each of the cell lines (Figure 4A). One fusion, RP11-96H19.1-RP11-446N19.1, was seen across all five clusters. It is not present in CCLE and is consistent with run-through transcription with constituent genes 264 kbp apart in the reference genome (Figure 4B). Some fusions were detected in the wrong cell line cluster (Figure 4A) and we hypothesize that long read sequencing errors in the cell barcodes have led to misassignment of reads in these cases. However, despite errors, these results demonstrate that JAFFAL enables fusions to be detected at the level of individual cells.

**Figure 4:**
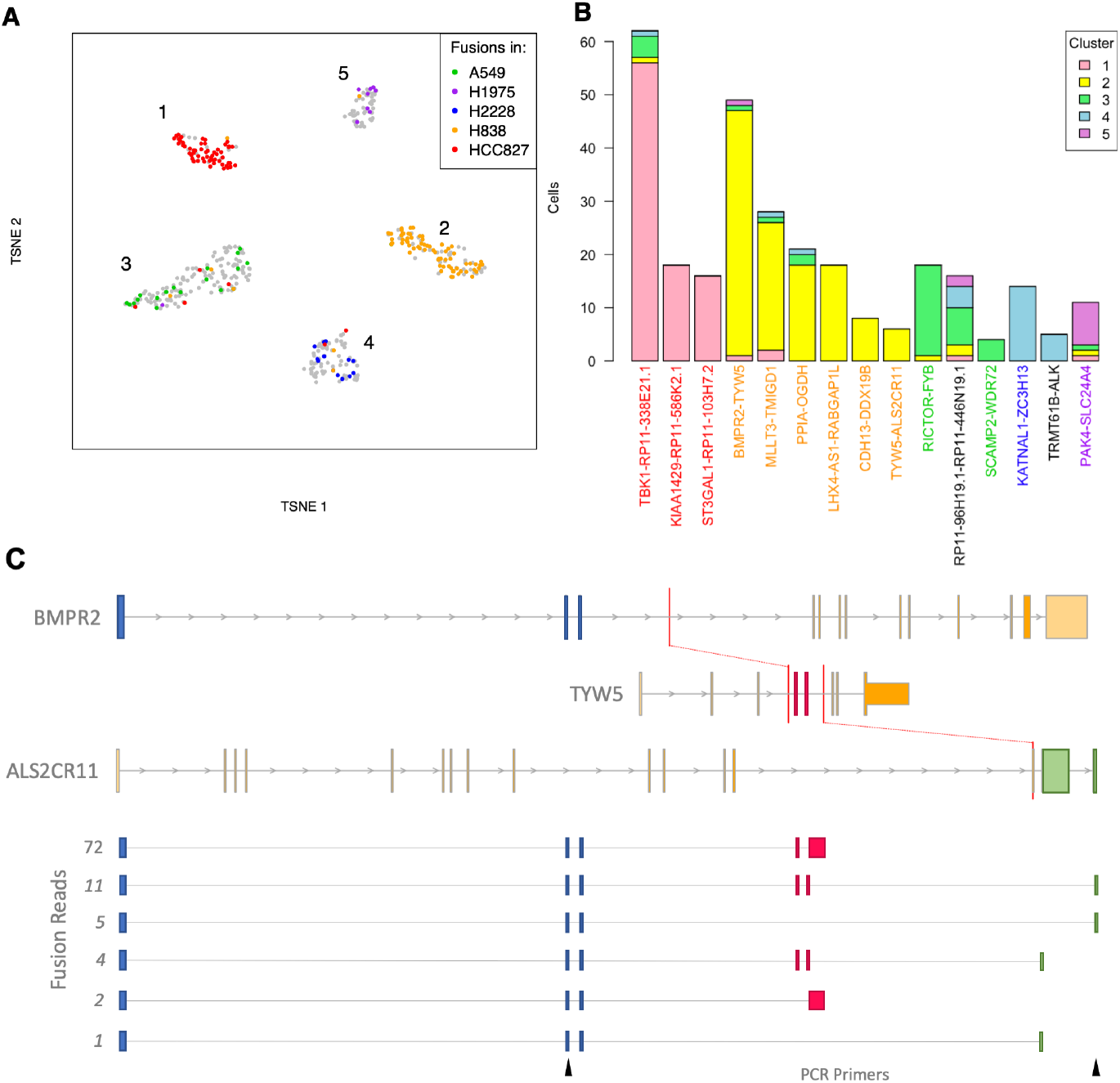
Detection of fusions in single cell ONT sequencing of five cell lines. A) t-SNE plot generated from short read gene expression. Colour indicates the cell line that a fusion detection is known to be in from CCLE. Grey indicates a cell with no detected CCLE fusion. B) For each of the 16 fusions detected by JAFFAL the number of cells identified in each of the five clusters is shown. Fusion labels are coloured according to the CCLE cell line they were previously identified in. Black indicates a novel fusion. C) JAFFAL identified BMPR2-TYW5 and TYW5-ALS2CR11 in the H838 cell line as belonging to the same transcript and forming the three-gene fusion BMPR2-TYW5-ALS2CR11 identified in 15 reads (two different isoforms). Expressed exons in the fusion transcript are shown in blue, red and green, with colour indicating the gene of origin. Red bars show the position of translocations seen in short read whole genome sequencing of H838 in CCLE. The breakpoint within ALS2CR11 falls within its third final exon and this exon appears to be spliced out. The six isoforms we identified for BMPR2-TYW5-ALS2CR11 and the number of long reads supporting each are also shown. The location of PCR forward and reverse primers which validated the translocation between BMPR2 and ALS2CR11 are shown in black (bottom).

### JAFFAL detects three-gene fusions

Recent analysis of rearrangements leading to fusions has described “bridged” fusions where genes are brought together through complex structural events that involve more than two genomic regions [15,43]. Although sequence from the bridged region is generally not transcribed, at least one instance of a three-gene fusion transcript has been reported [54]. Short read sequencing has limited the detection of three (or more) gene fusions as breakpoints often can not be linked within a fragment and short read fusion finding algorithms generally do not attempt to link breakpoints. Full transcript sequencing with long reads and new analysis algorithms can automatically discover these complex, linked events.

JAFFAL takes advantage of long reads to search for multiple fusion breakpoints within individual reads. We searched for multi-fusion reads across all our validation data and identified 14 three-gene fusions (Supplementary Table 9) in our PacBio, SGNex, patient data and single cell cell line datasets, with the majority, 9, from the highly rearranged cell line MCF-7. Four of the three-gene fusions had both their constituent breakpoints classed as High Confidence and the individual breakpoints were also seen in orthogonal data from short read sequencing (Supplementary Table 4). Interestingly, in all cases, a two-gene transcript which excluded one of the constituent fusions was also expressed and at a higher level than the three-gene isoform (Supplementary Table 4).

One of the high confidence three-gene fusions found by JAFFAL was BMPR2-TYW5-ALS2CR11 in single cell sequencing of the H838 cell line. It results from a complex rearrangement of a 2.5Mbp region on chromosome 2 and is supported by translocations found in CCLE whole genome sequencing [53] (Figure 4C). Long reads linked the BMPR2-TYW5 and TYW5-ALS2CR11 breakpoints in 6 cells. In 46 cells an alternative truncated transcript was also seen which links the BMPR2-TYW5 breakpoint to a novel exon extension event in TYW5 (Figure 4C). In both instances, the BMPR2-TYW5 breakpoint and second event were separated by 184 bp in the RNA. Although these transcripts could in theory be inferred with pair-end short read data, the linked events could not be covered by a single read of conventional length (150bp or less). In total we identified 6 distinct isoforms of the BMPR2-TYW5-ALS2CR11 fusion gene (Figure 4C), including transcripts where TYW5 is spliced out. The three gene fusion transcript BMPR2-TYW5-ALS2CR11 and its two gene transcript, BMPR2-ALS2CR11, were validated in the H838 cell line with PCR and Sanger sequencing (Supplementary Figure 6). This example illustrates that fusion finding with long reads can identify complex fusion transcripts which goes beyond just breakpoint discovery. For the first time we now have the tools to discover multi-rearranged genes and their alternative splicing in individual cells.

### Computational Resources

The computational resources required for JAFFAL and LongGF were benchmarked on a machine with 32 cores and 190GB of available memory. JAFFAL and minimap2 were given a maximum of four threads. LongGF which is single threaded, used one. JAFFAL completed in less than six hours and 21GB of memory on each of the nine healthy and cancer cell line bulk datasets described previously (Table 4). Despite running on only a single thread, LongGF used considerably less computational resources than JAFFAL. However, LongGF required reads which had already been mapped to the genome. Genome alignment using minimap2 was slower than JAFFAL, but required approximately the same memory. These results indicate that fusion calling on large long read sequencing cohorts is unlikely to be hindered by computational limitations using either fusion finder.

**Table 4:** Average and range (in parentheses) of run-time and memory consumed on nine benchmarking datasets by JAFFAL and LongGF.

|  |  | Run-time (hours) | Memory Consumption (GB) |
| --- | --- | --- | --- |
| JAFFAL (4 threads) |  | 2.6 (0.08 - 5.9) | 20.0 (19.8 - 21.1) |
| LongGF | Genome Mapping (4 threads) | 9.5 (0.1 - 21.2) | 22.6 (20.2 - 24.7) |
|  | LongGF (1 thread) | 0.4 (0.01 - 1.1) | 6.4 (0.8 - 13.3) |

## Conclusions

Long read sequencing is growing in popularity due to its ability to measure long stretches of sequence. A natural application is therefore detecting structural rearrangements, which in the transcriptome, can arise as fusions. However, very few computational methods exist for fusion detection from long read transcriptome sequencing. Here we introduce JAFFAL which is one of the first long read fusion finders. We demonstrated that JAFFAL is sensitive on simulated data over a range of read identities and coverage levels designed to mimic ONT and PacBio data. On real data, JAFFAL detected previously known fusions in cancer cell lines and patient samples with few false positives.

The ranking of fusions for prioritisation is an important feature of fusion finders. While alternative methods rely on the number of read support only, we demonstrate that other heuristics are powerful for separating artifacts from true fusions. By applying JAFFAL to samples sequenced with both direct RNA and cDNA we found a high rate of chimeric artifacts introduced during reverse transcription of libraries. We showed that these can be controlled by either sequencing RNA directly or by downranking fusions if their breakpoint does not coincide with exon boundaries.

Although the idea of using breakpoint positions as a heuristic in fusion ranking was first introduced for short read data [24], errors in long reads make precise breakpoints difficult to determine. JAFFAL overcomes this challenge by clustering reads into breakpoints and anchoring them to exon boundaries or the position with the maximum read support, per cluster.

This approach is one clear advantage of JAFFAL and was found to give fewer false positives compared to the competing long read tool, LongGF on cancer cell line data.

JAFFAL and LongGF were found to identify different fusions when applied to the MCF-7 cell line. Differences between fusions called by different tools on short read data are well documented, and this is why clinical pipelines often employ an ensemble approach combining the results of several fusion finding tools together, to identify actionable fusions [5,49,55]. It is likely that long read fusion finding will also benefit from multiple methods being available, and JAFFAL represents an important early contribution towards this.

A limitation of JAFFAL is its dependence on annotated transcripts. Fusions which incorporate intergenic or intronic sequences at a breakpoint are not detected. Hence complex fusions such as IGH-CRLF2 in our patient sample will be missed. This highlights an area for further development in long read fusion finding. As shown in our simulation, the detection of fusions is limited by their coverage, which is directly related to expression levels, and by error rates in the data. However, sequencing accuracy from long read technologies is improving and is likely to benefit fusion finding with JAFFAL in the future.

Finally, long read sequencing has a number of novel advantages over short reads. An exciting development has been the use of long reads in conjunction with single cell RNA sequencing, which enables the full transcriptomes of individual cells to be sequenced. Here we demonstrate that fusions can be called in this data, adding an extra modality to single cell analysis, providing many new opportunities to study the heterogeneity of tumours. Long reads enable novel events to be linked over the full length of fusion transcripts meaning additional variants, such SNPs, splicing or other fusions can be phased. JAFFAL thus allows the automatic detection of three gene fusions and we demonstrated the detection of a novel three gene event, BMPR2-TYW5-ALS2CR11 in the lung cancer cell line H838. ONT sequencing has several further advantages including the profiling of the epitranscriptome and rapid and remote sequencing. Combined with fusion finding, these technological advances have the potential to enable greater understanding of the mechanisms driving tumours and the potential to bring clinical diagnostics to remote areas.

## Materials and Methods

### JAFFAL pipeline

JAFFAL is a multi-stage bpipe [27] pipeline for fusion detection. A brief outline of its steps follow. Fastq files are unzipped and converted to fasta prior to alignment to the human reference transcriptome, gencode version 22 for hg38, with minimap2 version 2.17 and option -*x map-ont*. Alignments to the transcriptome are then processed with a custom C++ program, which identifies reads which align to two distinct genes. The two alignment intervals within a read must have no more than 15bp of overlap, no more than a 15bp gap and be on the same strand. Fusion candidate reads are then extracted into a fasta file and aligned to the reference genome, hg38 using minimap2 with option -*x splice*. Genome alignments are processed using a custom R script. It first finds the breakpoint positions in the genome and filters any where the start and end are within 10kbp of each other in an order consistent with regular transcription. Next alignments are compared against annotated transcripts, and breakpoints realigned to exon boundaries as described in Results. Fusions involving the mitochondrial chromosome are filtered out (by default). Reads are then aggregated by breakpoint, and clustered using the following algorithm:

~~~
For each fusion gene:
    For each breakpoint:
    Identify the next closest breakpoint in euclidean distance
    If (breakpoint is within an exon) and (next closest breakpoint < 50bp ):
           flag breakpoint for reassignment
    Order flagged breakpoints from lowest to highest supporting reads
    For each flagged breakpoint:
           Add the number of supporting reads to the next closest breakpoint
           Remove breakpoint
~~~

Next, breakpoints are classified as either high confidence, low confidence, potential trans-splicing (Figure 1) or run-through transcription. Run-through transcription includes any fusion with breakpoints that are within 200kbp of each other in an order consistent with regular transcription and these are filtered out by default. Information on whether the fusion is in frame and seen in the Mitelman database is added. Breakpoints are then reported in a csv output file. Finally, all candidate fusion reads are compared to the final fusions gene list. Reads consistent with multiple fusions are aggregated and reported. The code for JAFFAL is open source and available at https://github.com/Oshlack/JAFFA. The results presented in this manuscript were generated with JAFFAL version 2.2 run with the flag -*n4* (4 threads).

### LongGF

Samples were first mapped to version hg38 of the human reference genome using minimap2 version 2.17 with flags *-t4* and *-ax splice*. Mapped reads were name sorted with samtools before being processed with LongGF version 0.1.1. We ran LongGF with the annotation file gencode.v22.chr_patch_hapl_scaff.annotation.gtf downloaded from https://www.gencodegenes.org/human/release_22.html. This annotation used the same gene names as the reference provided to JAFFAL. We used options “*100 50 100 0 0 1*” which were recommended apart from the number of reads support which we lowered from 2 to 1 to assess sensitivity. Fusions involving a gene on the mitochondrial chromosome were removed to allow consistent comparison against JAFFAL which removes these by default.

### Simulation

Simulated fusion transcripts created by Haas et al. [10] were downloaded and split into 25 fasta files for each of the 25 combinations of coverage-levels (1,2,10,50,100) and read identities (75%, 80%, 85%, 90%, 95%). Sequencing reads were then simulated using Badread version 0.1.5 with corresponding coverage and read identity levels set through the parameters --*quantity* <*coverage*> and --*identity* <*read identity*>,*95,5* respectively. The error model was set to either pacbio or nanopore with the parameters --*error_model* and --*qscore_model*. To simplify the simulation, we switched off artifacts with the options --*junk_reads 0* --*random_reads 0* --*chimeras 0*. Chimeras in long reads were assessed with real data rather than simulation.

### Comparison

For the simulation, fusions were classed as true positives if there was a simulated breakpoint within 1kbp euclidean distance of the reported breakpoint. For cell lines data we matched fusions to those previously validated (Supplementary Table 2) and between fusion finders using gene names. Novel breakpoints within known fusion gene pairs were considered true positives. Unless stated otherwise, comparisons were performed at the fusion gene-level, meaning fusions with multiple breakpoints were counted as a single true or false positive, and given the ranking of their highest ranked breakpoint. LongGF does not report fusions in transcriptional order, hence if a known fusion gene pair was not seen, we also checked the reciprocal gene order and counted these as a match if found. Similarly, for known fusions involving antisense genes we counted the sense gene name as a match if reported.

### Patient Sequencing

For the B-ALL patient sample, an ONT sequencing library was generated from approximately 100 ng of total RNA using the ONT cDNA-PCR Sequencing Kit (SQK-PCS109) and sequenced using a MinION Nanopore sequencer on a R9.4 flow cell (FLO-MIN106). Basecalling was performed using Guppy version 4.2.3.

### Single cell analysis

Cellular barcodes were annotated to long reads using FLAMES [20]. JAFFAL was then run on pooled reads. A custom script, get_cell_barcodes_by_fusion.bash, which is available in JAFFA, was used to generate a table of fusions by cell barcode. Only fusions classes as high confidence and found in four or more cells were analysed further. Matched short read gene expression count data from Tian et al. [20] was downloaded from https://github.com/LuyiTian/FLTseq_data/blob/master/data/PromethION_scmixology1.zip and analysed with Seurat [56]. A list of fusions called in short read CCLE data was obtained from the CCLE data portal, https://portals.broadinstitute.org/ccle.

### Validation of BMPR2-TYW5-ALS2CR11 in the H838 cell line

RNA was extracted from H838 and HEK293T cells using NuceloZOL (Macherey-Nagel), followed by cDNA synthesis using SuperScript III (Invitrogen) with OligoDT or random hexamer primers. PCR reactions were performed using Q5® High-Fidelity DNA Polymerase (NEB) with the following primers: *BMPR2 Fusion F:* GGTAGCACCTGCTATGGCCT; *BMPR2 Fusion R:* CTAAGCCTGATGAAACCATTCGACG; *GAPDH F:* TGAAGGTCGGAGTCAACGGATTTGGT; *GAPDH R:* CATGTGGGCCATGAGGTCCACCAC. BMPR2 PCR products amplified from the H838 cDNA were purified from agarose gel using the NucleoSpin Gel and PCR Clean-up kit (Macherey-Nagel), and subjected to sanger sequencing using the BMPR2 primers listed above.

## Abbreviations

ONT: Oxford Nanopore Technologies
ALL: acute lymphoblastic leukemia
AML: acute myeloid leukemia
SRA: Sequence Read Archive
CCLE: Cancer Cell Line Encyclopedia
PCAWG: Pan-Cancer Analysis of Whole Genomes
TCGA: The Cancer Genome Atlas

## Declarations

### Ethics approval and consent to participate

ONT sequencing of the B-ALL patient sample was conducted under Peter MacCallum Cancer Centre Ethics Committee approval (HREC/17/PMCC/163) and all experiments were performed in accordance with the Declaration of Helsinki.

### Consent for publication

Not applicable

### Data Availability

Simulated data is available for download from figshare [57]. NA12878 cell line data was downloaded from https://github.com/nanopore-wgs-consortium/NA12878. SGNex ONT cell line sequencing is available at https://github.com/GoekeLab/sg-nex-data. PacBio sequencing of MCF-7, HCT-116 and SK-BR-3 cell lines are available from the sequence read archive (SRA) under accessions SRP055913, SRP091981 and SRP150606 respectively. The patient sample from Lui et al. [26] is available from SRA under accession SRP267910. The long read data for the B-ALL patient sample is available from the corresponding authors on reasonable request. The ONT single cell sequencing is available under SRA accession SRP273167.

### Authors’ contributions

NMD and AO conceived of ideas in this manuscript and designed the research. NMD wrote the software, performed all the analysis and wrote the preliminary draft of the manuscript. JG and YC provided cancer cell line data and contributed to concept development. GLR and PB provided patient data. TS and PGE performed fusion validation. NMD, GLR, TS, PB, PGE, JG and AO revised the manuscript. All authors read and approved the final manuscript.

## Acknowledgements

The authors would like to thank Mike Clark and Ricardo De Paoli-Iseppi (Centre for Stem Cell Systems, Department of Anatomy and Neuroscience, The University of Melbourne) for their assistance with experimental design, library preparation and sequencing of patient samples. We would also like to thank Matt Ritchie and Luyi Tian for access to long read single cell datasets. Computing resources for this work were provided by PeterMac and The University of Melbourne Science IT. P.G.E. acknowledges the support of the Samuel Nissen Charitable Foundation.

