## Supplementary Figures 1-6 and Supplementary Tables 2,3,7 and 9 for "JAFFAL: Detecting fusion genes with long read transcriptome sequencing"

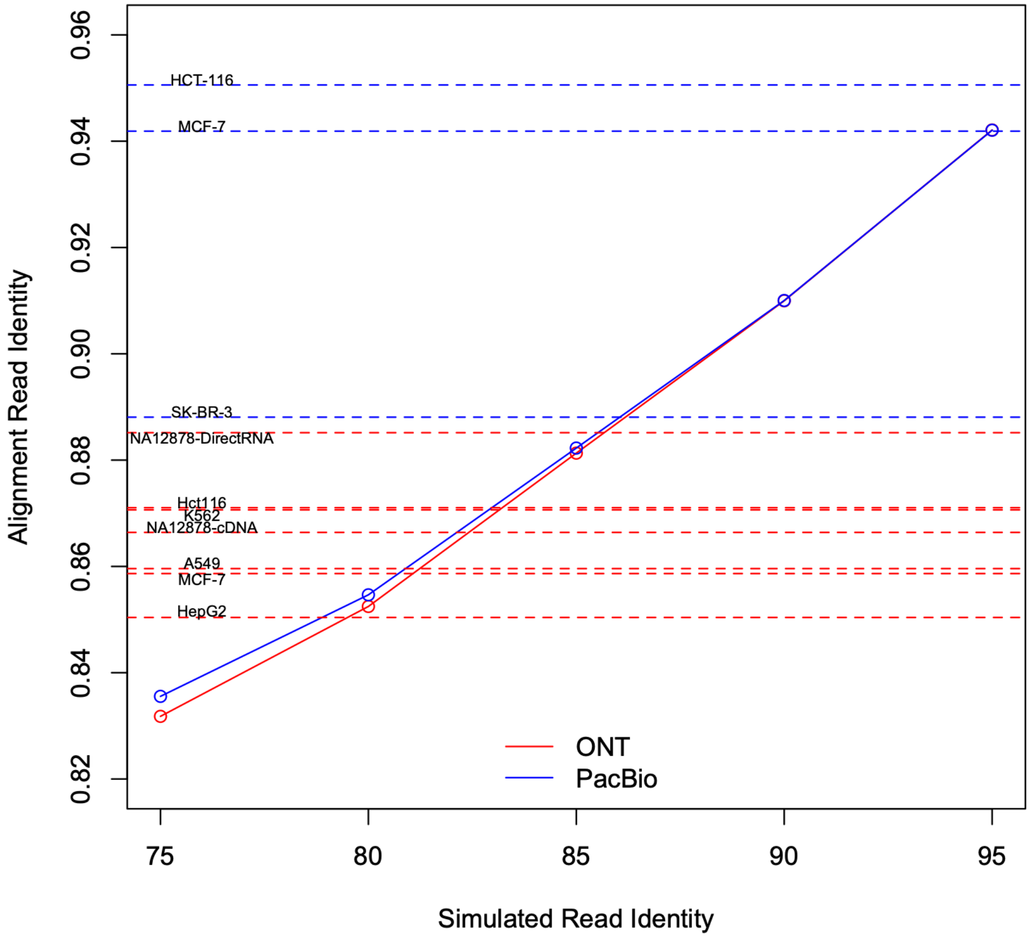


**Supplementary Figure 1: Simulated and aligned read identifies for the datasets used to evaluate JAFFAL's performance**. Solid curves show the simulated read identities for the artificially generated data as provided to badread and their corresponding read identities as measured from alignment to the reference transcriptome with minimap2. The sequence identity is defined by matched bases divided by alignment length. The alignment read identity is generally higher than that simulated because successful alignment is biased towards low error sequence. The horizontal dashed levels correspond to the alignment read identities measured in the real cell line datasets described in the manuscript.

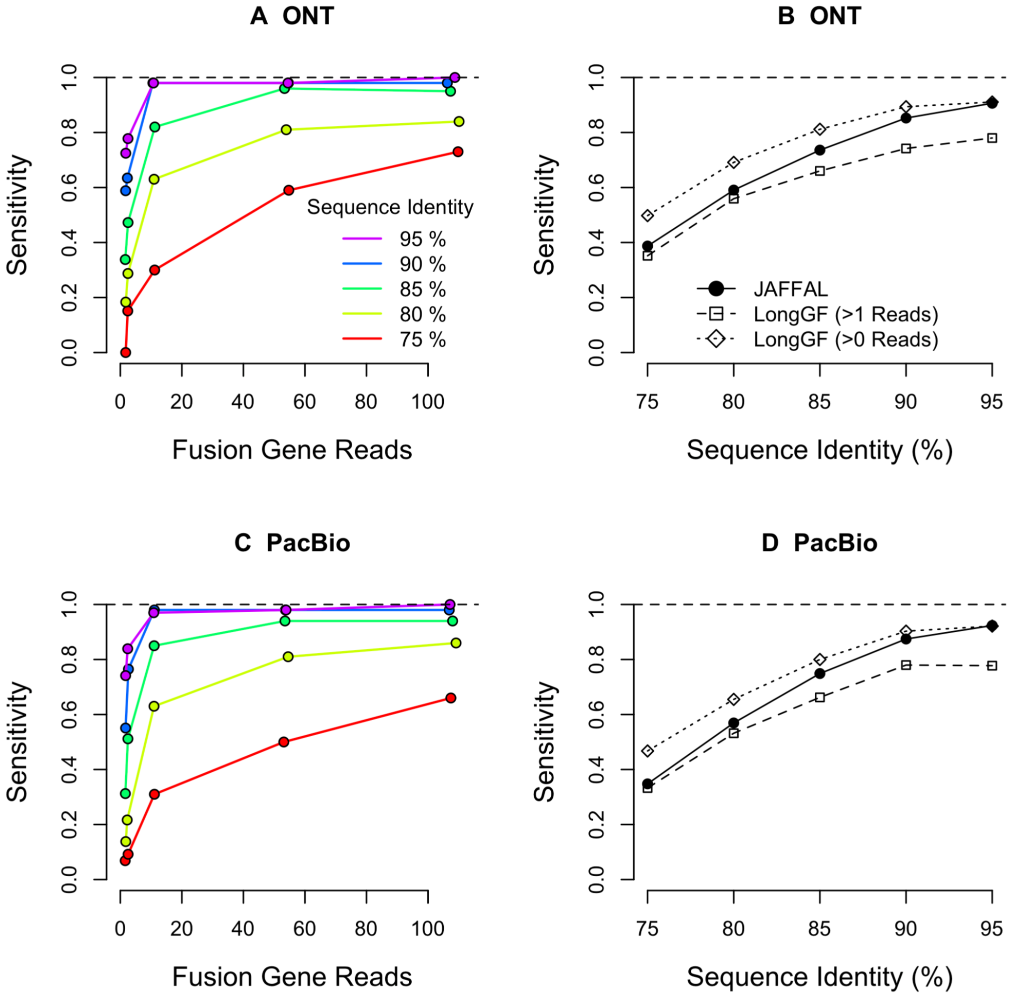


**Supplementary Figure 2: Fusion finding sensitivity on ONT and PacBio simulated data without background**. A) and C) The fraction of simulated fusions detected (y-axis) by JAFFAL across a range of fusion coverage levels (x-axis) and read identity levels (red-purple). B) and D) The fraction of simulated fusions detected (y-axis) by JAFFAL and LongGF for sequence identity levels of 75-95%. ONT and PacBio data were simulated using the same fusion sequence, coverage level and read identity parameters. JAFFAL detected more fusions than LongGF when LongGF was run with default parameters (>1 read support), but fewer when LongGF was allowed to report fusions with just one read support. This intermediate behaviour of JAFFAL is consistent with its reporting fusions with one read support conditional on the breakpoint coinciding with exon boundaries. At low read identities, this condition is more likely to fail due to poor alignment. LongGF reported fewer simulated fusions when background reads were included in the data (Manuscript Figure 2B).

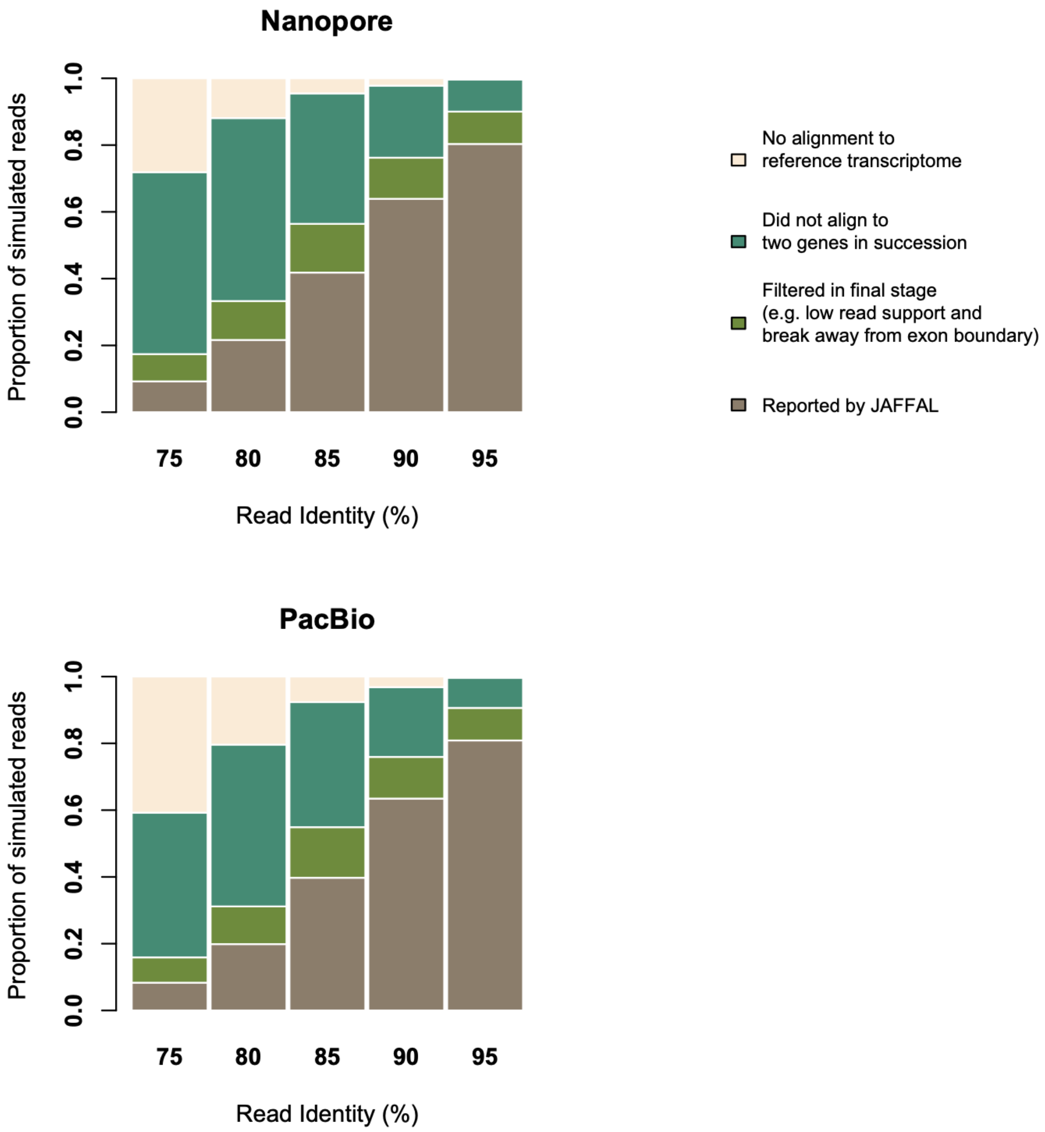


**Supplementary Figure 3: Proportion of simulated fusion reads lost in various stages of the JAFFAL pipeline**. Fusion reads are predominantly lost due to failure to align to the reference transcriptome (cream and dark green). As read identity increases alignment becomes more accurate and a greater proportion of reads are identified by JAFFAL (brown).

| | | cRNA - Raw | cDNA - PoreChop |
|---|---|---|---|
| Total Reads Processed | | 25,418,307 | 25,286,945 |
| Fusion genes called by JAFFAL | High Confidence | **8** | **9** |
| | Low Confidence | 94 | 100 |
| | Potential Trans-splicing | 412 | 410 |

**Supplementary Table 2: The number of fusions called in the non-cancer cell line NA12878 from ONT amplified cDNA before and after processing the data with PoreChop.** Two full transcripts including adapters may be sequenced in succession in a single ONT read. To examine the impact of this type of chimera on fusion calling, we applied PoreChop (https://github.com/rrwick/Porechop) to data from NA12878. PoreChop searches and removes adapter sequences and splits reads where internal adapters are found. Approximately 30,000 reads were split by PoreChop. The number of fusions called by JAFFAL remained similar after running PoreChop.

| | | Direct RNA | cDNA downsamples |
|---|---|---|---|
| Total Reads Processed | | 14,971,421 | 14,971,421 |
| Fusion genes called by JAFFAL | High Confidence | **4** | **7** |
| | Low Confidence | 5 | 43 |
| | Potential Trans-splicing | 344 | 249 |

**Supplementary Table 3: The number of fusion genes called in NA12878 from ONT direct and amplified cDNA downsampled to the same number of reads.** The cDNA sample retains significantly more low confidence fusion calls than the direct RNA sequence even after downsampling to the same number of reads.

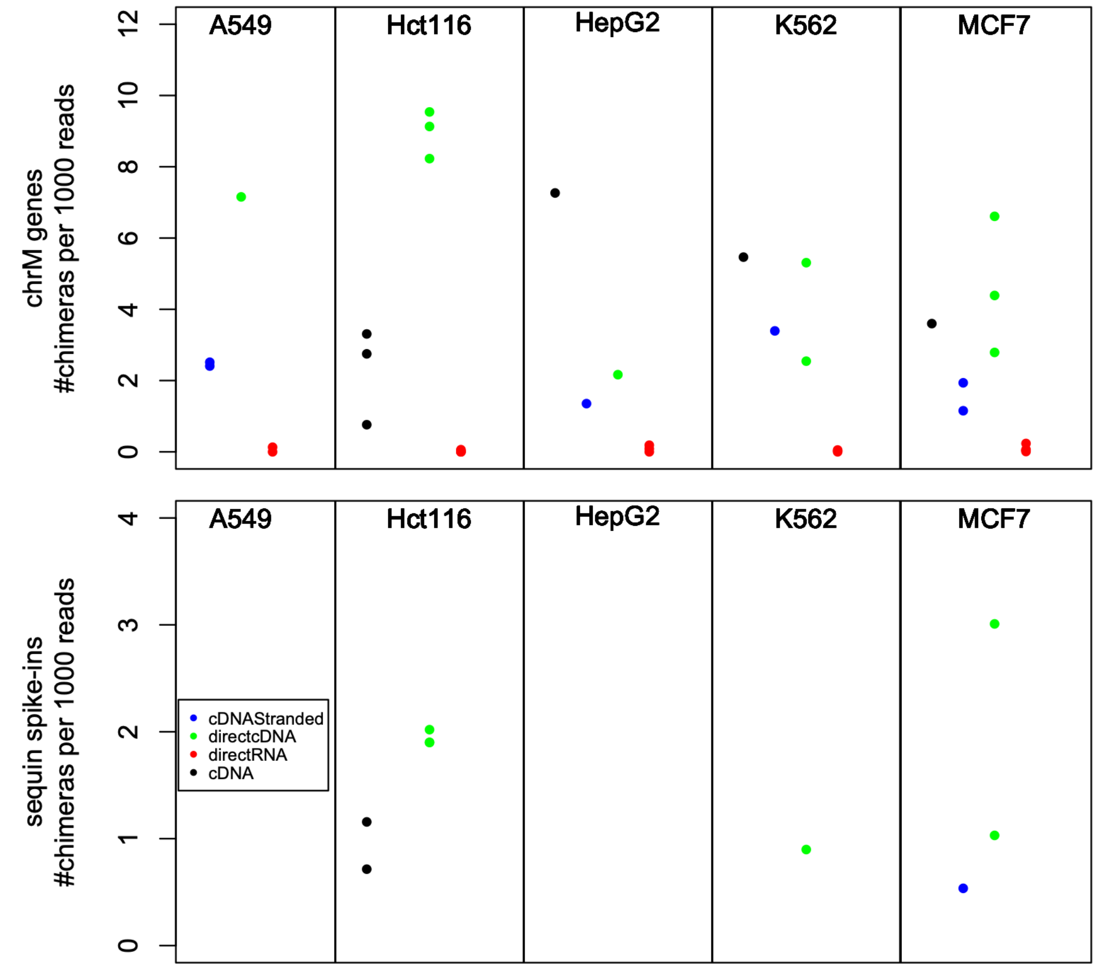


**Supplementary Figure 4. Estimated rates of chimeric artifacts in the SGNex dataset.** SGNex data consists of multiple replicates sequenced using different ONT protocols: direct RNA, direct cDNA and amplified cDNA for five cell lines. Two types of known false positives were used as indicators of chimeric artifacts, (top) fusions between a gene on the mitochondrial chromosome and a gene not on the mitochondrial chromosome, and (bottom) fusions between Sequin spike-in transcripts and regular genomic genes, in samples where Sequin spike-ins were added. JAFFAL's default filtering of mitochondrial genes was switched off and its reference supplemented with Sequin sequences. JAFFAL was run on each replicate, and initial candidate fusion reads (ie. those identified after reference transcriptome alignment but before reference genome alignment), were used to calculate the number of chimeric reads. This was then normalised by the total number of reads aligning to either mitochondrial (top) or Sequin (bottom) genes.

| Fusion | Tool | Predicted Breakpoint | Reads Support | Fusion Rank |
|---|---|---|---|---|
| BCR-ABL1 | JAFFAL | chr22:23,182,239 - chr9:130,854,064 | 5 | 5th |
| | LongGF | chr22:23,182,237 - chr9:130,854,060 | 5 | 8th |
| RUNX1-RUNX1T1 | JAFFAL | chr21:34,859,474 - chr8:92,017,363 | 8 | 1st |
| | LongGF | chr21:34,859,474 - chr8:92,017,365 | 11 | 2nd |
| IGH-CRLF2 | JAFFAL | Not detected | - | - |
| | LongGF | Not detected | - | - |

**Supplementary Table 7:** Clinically relevant fusions detected in two patient samples by JAFFAL and LongGF. JAFFAL and LongGF show similar fusion ranking and read support. JAFFAL detects the exact breakpoints known from short-read sequencing

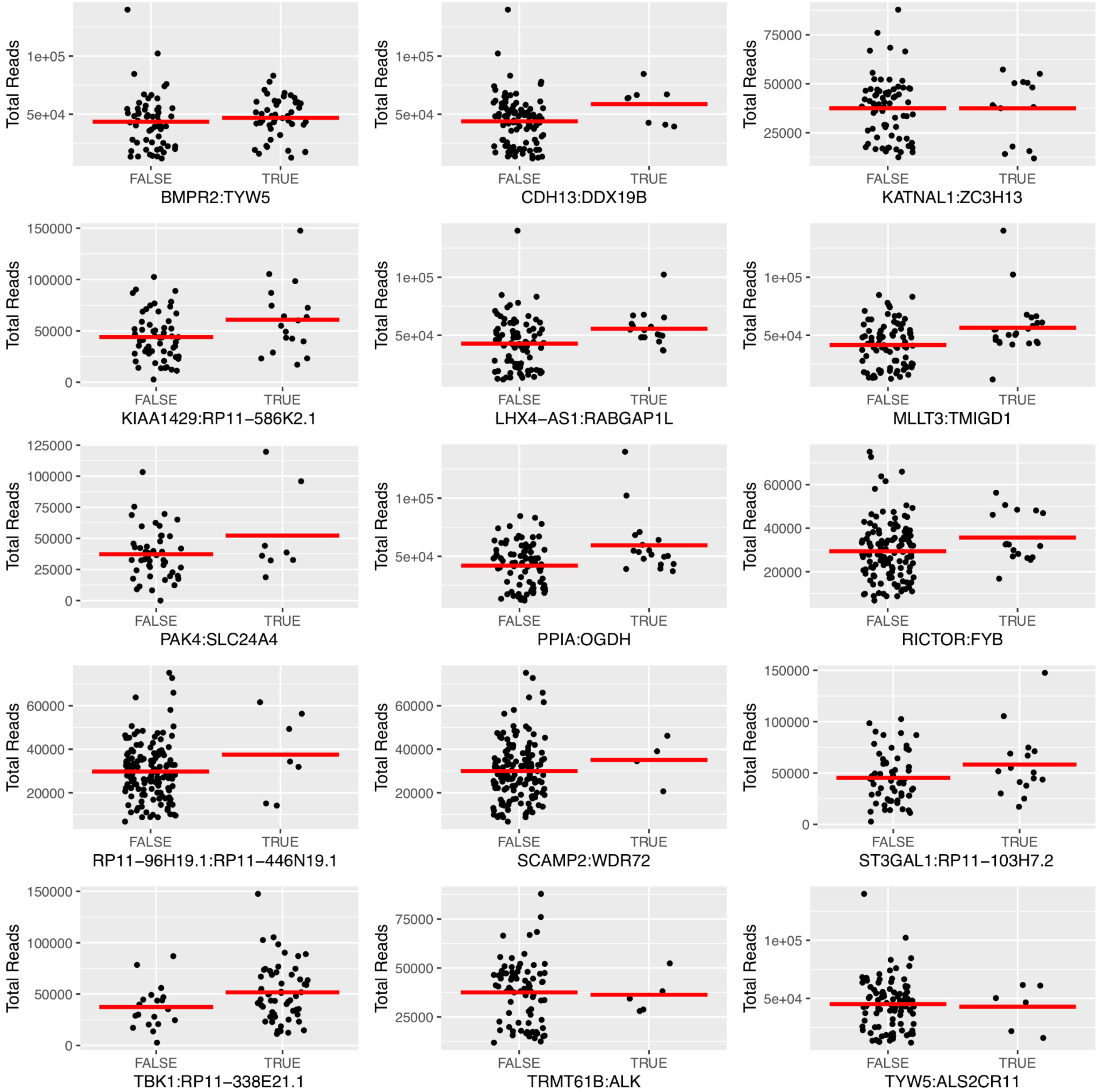


**Supplementary Figure 5: Library size of single cells sequencing where fusions were or were not identified.** For each fusion in the long read single cell sequencing data, we identified all cells in the corresponding gene expression cluster (see Manuscript). The total number of reads for each cell is shown (black) and mean (red bar) for cells where the given fusion was either identified (TRUE) or not (FALSE). Whether fusions could be identified in individual cells is likely to be a combination of total sequencing depth for the cell, heterogeneity in gene expression of the fusion and sampling. For most cells only a single fusion read was detected.

| Cell line | Dataset | Fusion | Reads | Breakpoint Read | Breakpoint Classes | Orthogonal Evidence |
|---|---|---|---|---|---|---|
| H838 | scRNA-Seq | BMPR2:TYW5:ALS2CR11 | 15 | 89:15 | High:High | Both fusions seen in RNA and WGS from CCLE (Barretina et al, Nature, 2012) |
| H2228 | scRNA-Seq | RP11-448A19.1:SND1:CFTR | 4 | 18:4 | Low:High | Both fusions seen in RNA from CCLE |
| MCF7 | PacBio | TXLNG:SYAP1:RRM2 | 4 | 29:6 | High:Low | - |
| H838 | scRNA-Seq | XPR1:LHX4-AS1:RABGAP1L | 2 | 2:64 | High:High | - |
| MCF7 | PacBio | BCAS4:BCAS3:REG4 | 2 | 1304:2 | High:High | Both fusions seen in RNA from ENCODE (Davidson et al., Genome Med. 2014) |
| MCF7 | SGNex | GBF1:MACROD2:C14orf132 | 1 | 13:2 | High:High | Both fusions seen in matched Illumina data from SGNex |
| H838 | scRNA-Seq | TRIP13:BMPR2:TYW5 | 1 | 1:89 | TransSplicing:High | - |
| MCF7 | SGNex | YY1:PPP1R12A:EVL | 1 | 1:1 | TransSplicing:TransSplicing | - |
| MCF7 | SGNex | VMP1:BTBD1:YPEL5 | 1 | 1:1 | TransSplicing:TransSplicing | - |
| MCF7 | SGNex | RAD51B:CCDC170:EPB41L5 | 1 | 1:1 | TransSplicing:TransSplicing | - |
| K562 | SGNex | MPV17:TCERG1:CREBZF | 1 | 1:1 | TransSplicing:TransSplicing | - |
| MCF7 | SGNex | IKZF2:NCOR1:SPATA33 | 1 | 1:1 | TransSplicing:TransSplicing | - |
| MCF7 | SGNex | CFL1:SLC4A7:URI1 | 1 | 1:1 | TransSplicing:TransSplicing | - |
| MCF7 | PacBio | COPS7B:AVL9:ZFYVE1 | 1 | 1:1 | TransSplicing:TransSplicing | - |

**Supplementary Table 9: Three-gene fusions identified by JAFFAL on cell line validation datasets.** Reads indicated how many reads spanned the three genes. Breakpoint reads give the number of reads reported by JAFFAL for each individual fusion (separated by ":" in gene order). Breakpoint Classes indicated the classification of the individual fusions, as reported by JAFFAL.

(A)

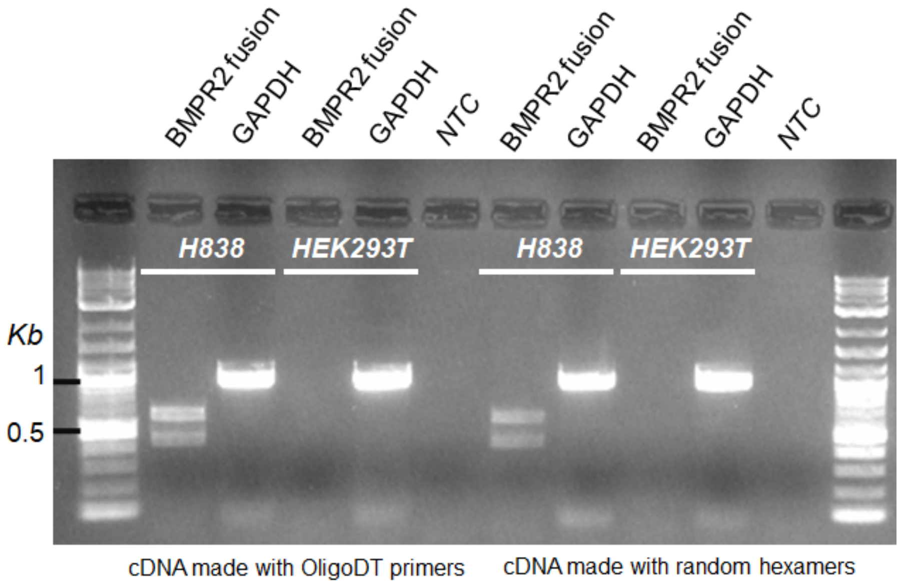


(B)

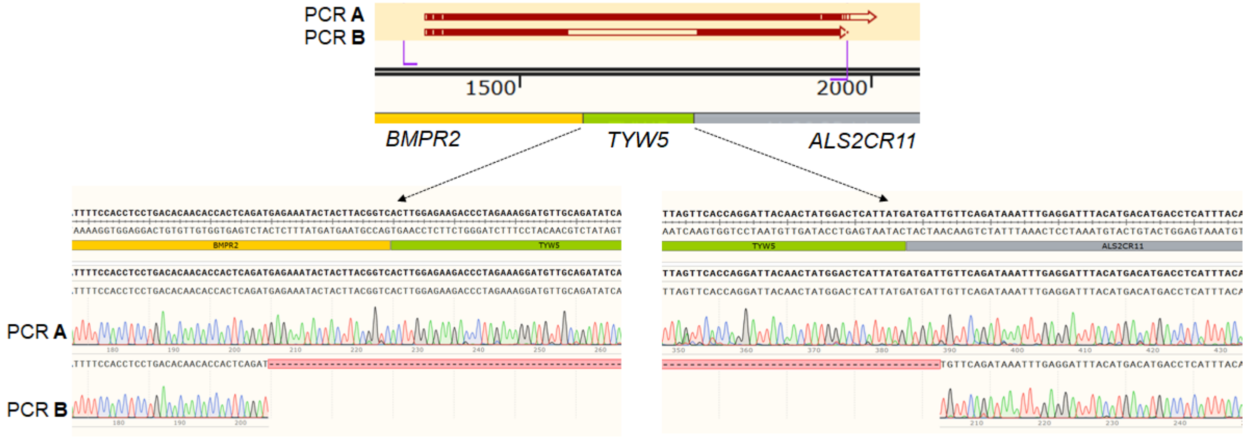


**Supplementary Figure 6: Validation of BMPR2-TYW5-ALS2CR11 fusion**. (A) PCR validation of BMPR2-TYW5-ALS2CR11 fusion in cDNA from H838 cells synthesized with OligoDT primers or random hexamers. cDNA from HEK293T cells was used as a negative control. (B) Sanger sequencing of the top band (PCR A) and lower band (PCR B) further confirmed these correspond to the three gene fusion BMPR2-TYW5-ALS2CR11 and it's two gene transcript, BMPR2-ALS2CR11, respectively.
